# MetaboDirect: An Analytical Pipeline for the processing of FTICR-MS-based Metabolomics Data

**DOI:** 10.1101/2022.06.10.495699

**Authors:** Christian Ayala-Ortiz, Nathalia Graf-Grachet, Viviana Freire-Zapata, Jane Fudyma, Gina Hildebrand, Roya AminiTabrizi, Cristina Howard-Varona, Yuri E. Corilo, Nancy Hess, Melissa Duhaime, Matthew Sullivan, Malak Tfaily

## Abstract

**Background:** Microbiomes are now recognized as main drivers of ecosystem function ranging from the oceans and soils to humans and bioreactors. However, a grand challenge in microbiome science is to characterize and quantify the chemical currencies of organic matter (i.e. metabolites) that microbes respond to and alter. Critical to this has been the development of Fourier transform ion cyclotron resonance mass spectrometry (FTICR-MS), which has drastically increased molecular characterization of complex organic matter samples, but challenges users with hundreds of millions of data points where readily available, user-friendly, and customizable software tools are lacking.

**Results:** Here, we build on years of analytical experience with diverse sample types to develop MetaboDirect, an open-source, command-line based pipeline for the analysis, visualization, and presentation of metabolomics data by direct injection FTICR-MS after molecular formula assignment has been performed. When compared to all other available FTICR software, MetaboDirect is superior with respect to its compute time as it only requires a single line of code that launches a fully automated framework for the generation and visualization of a wide range of plots, with minimal coding experience required. Among the tools evaluated, MetaboDirect is also uniquely able to automatically generate biochemical transformation networks (*ab initio*) based on mass differences that provide a comprehensive experimental assessment of metabolite connectives within a given sample or a complex metabolic system, thereby providing important information about the nature of the samples and the set of the microbial reactions or pathways that gave rise to them. Finally, for more experienced users, MetaboDirect allows users to customize plots, outputs, and analyses.

**Conclusion:** Application of MetaboDirect to FTICR-MS-based metabolomics datasets from a marine phage-bacterial infection experiment and a *Sphagnum* leachate microbiome incubation experiment showcase the exploration capabilities of the pipeline that will enable the FTICR-MS research community to evaluate and interpret their data in greater depth and in less time. It will further advance our knowledge of how microbial communities influence and are influenced by the chemical makeup of the surrounding system. Source code and User’s guide of MetaboDirect are freely available through (https://github.com/Coayala/MetaboDirect) and (https://metabodirect.readthedocs.io/en/latest/) respectively.

## BACKGROUND

Microorganisms play crucial roles in a host of fundamental ecological processes and are needed for maintaining a healthy global ecosystem [1, 2]. They are responsible for the mobilization, transformation, and storage of natural organic matter (NOM)-the complex mixture of organic compounds present within any system-thus driving the cycling of elements essential for life (e.g., carbon, nitrogen, sulfur) [2-5]. Microorganisms further contribute to the NOM pool, especially the dissolved organic matter (DOM) pool, in both aquatic and terrestrial systems [3, 6]. Environmental conditions such as temperature, and water availability can strongly influence the microbial community structure and function and thus its interaction with NOM [7, 8].

Microorganisms are capable of directly assimilating low-molecular-weight-DOM (< 600 Da) for their metabolic processes, while also producing microbial-derived products and residues that become integral components of NOM such as proteins, polysaccharides, and cell wall polymers [9]. The quantity and the quality of OM in a given ecosystem are dependent on its microbiome composition and the environmental conditions present in that system. They are also dependent on several microbial regulatory processes, such as transcription, translation, protein interactions, and their interactions with the biotic and abiotic components of the system [10-14]. Characterizing OM molecular composition is therefore vital for understanding the role that microorganisms play in all major element biogeochemical cycles and can constitute an important predictor of the response of the biological systems to environmental perturbations [8, 15].

Natural organic matter is a complex mixture of organic compounds whose size and other molecular properties are perceived as a continuum [16, 17]. The resolution of the components of this mixture, require the use of the substance-specific, molecular-level order mass spectra residing in the (sub)millimass space (mDa), which cannot be accessed by low-resolution mass spectrometers [16]. However, recent advances in analytical mass spectrometry techniques and in particular the introduction of high-resolution mass spectrometry (HR-MS) have allowed organic compounds to be identified based on ultra-high mass accuracy, and has led to more sensitive, selective, robust, and repeatable analyses [16-18]. Thus, FTICR-MS has evolved during the past two decades into a powerful tool to study the molecular composition of small-molecule organic complex mixtures (e.g., DOM in ocean waters and soil organic matter (SOM)) in diverse ecosystems [18].

One analytical approach using FTICR-MS is direct injection mass spectrometry (DI-MS), which involves the introduction of liquid samples directly into the mass spectrometer without an attached fractionation step. This technique considerably reduces the analysis time, as it is amenable to the use of auto sample handlers allowing to process hundreds of samples per day. Even though DI-MS has ample coverage and can detect a wide range of compounds (e.g., lipids, sugars, amino acids or lignin), some drawbacks are its inability to separate chemical isomers, lack of fine resolving power and most importantly a limited molecular search space due to ion suppression [19]. Nonetheless, the use of direct infusion FTICR-MS (DI FTICR-MS, or just FTICR-MS) can provide a comprehensive overview of the molecular profile of the NOM and a baseline to understand how biological systems respond to changes in the biotic or abiotic factors acting upon them. A wide range of studies have used FTICR-MS as a powerful tool to characterize NOM changes in environmental samples [20-23] and its use continues to increase every day.

Numerous tools for signal processing and assignment of molecular formulas of raw FTICR-MS data exists, including proprietary tools but also open-source software such as Formularity [24]. Most recently CoreMS [25] provides a comprehensive software framework, including signal processing and sample agnostic molecular formula assignment. Signal processing and molecular formula assignment produce large data matrices containing the elemental composition and measured intensity of the peaks present in each sample. These large datasets often require specialized data analysis pipelines for filtering, interrogation, comparison, and visualization [26]. Open-source software and pipelines available for the analysis and visualization of FTICR-MS data include web-based applications such as UltraMassExplorer (UME) [27], FREDA [28], MetaboAnalyst [29], and DropMS [30]. Even though the graphic user interface (GUI) of these software packages is user friendly, these types of software can be very restrictive in their use and do not allow users to fully customize their analysis to their research needs. Other more programming-friendly software packages include the visualization toolss *i-*van Krevelen [31] and OpenVanKrevelen [32] and the more comprehensive data analysis tools PyKrev [33] and *ftmsRanalysis* [26]. These approaches however require users to be competent in coding and programming in either R (for ftmsRanalysis) or Python, which further limits their usefulness for researchers without those skills. Thus, despite the broad availability of software packages for the analysis of FTICR-MS data, they often incur in a compromise between flexibility/customizability and user-friendliness that we aim to address with MetaboDirect by making it a fully automated pipeline capable of easily generating all of the figures, plots and analysis that are commonly used by the scientific community to visualize, analyze, and interpret FTICR-MS datasets (Table 1).

**Table 1.**
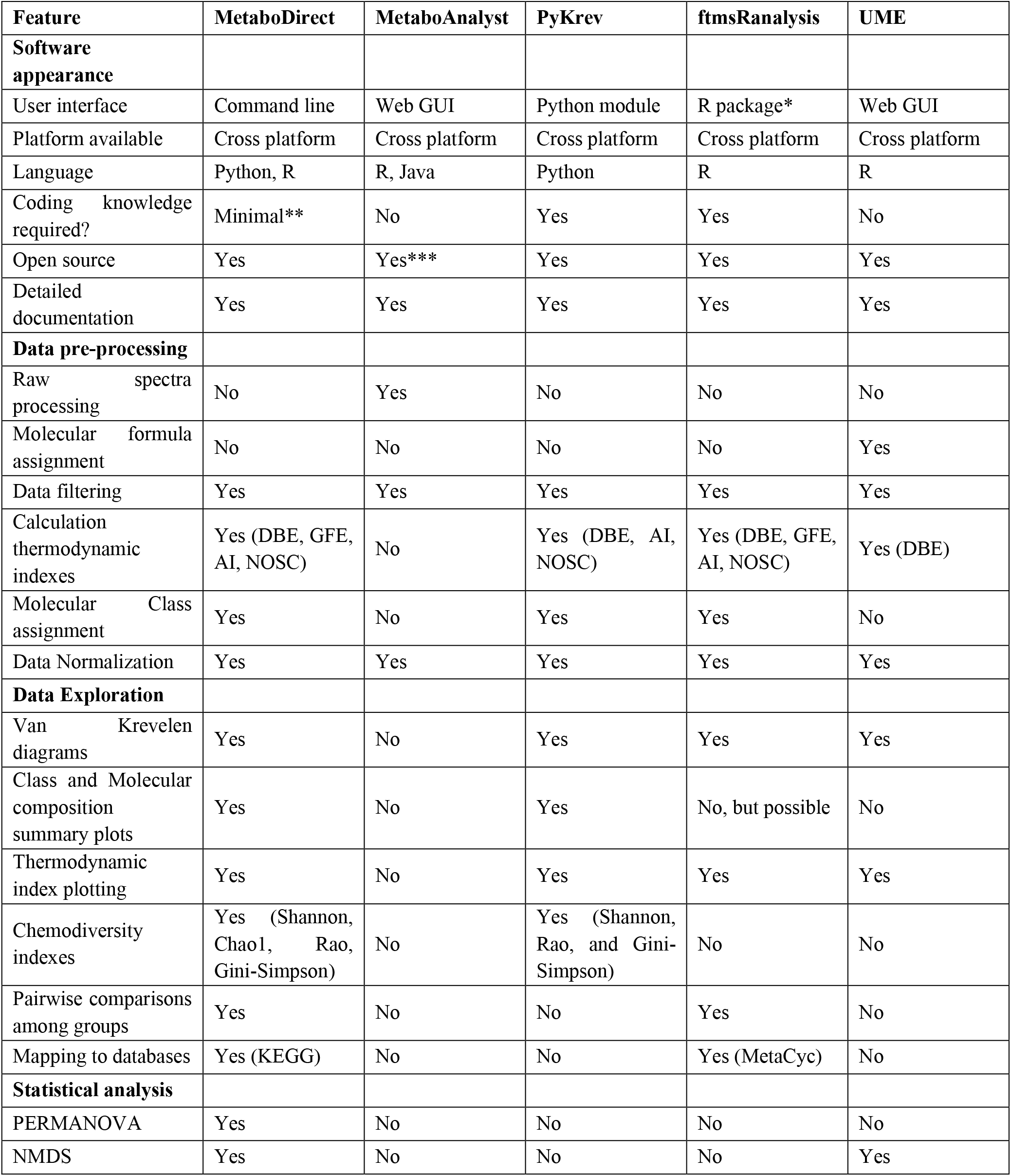
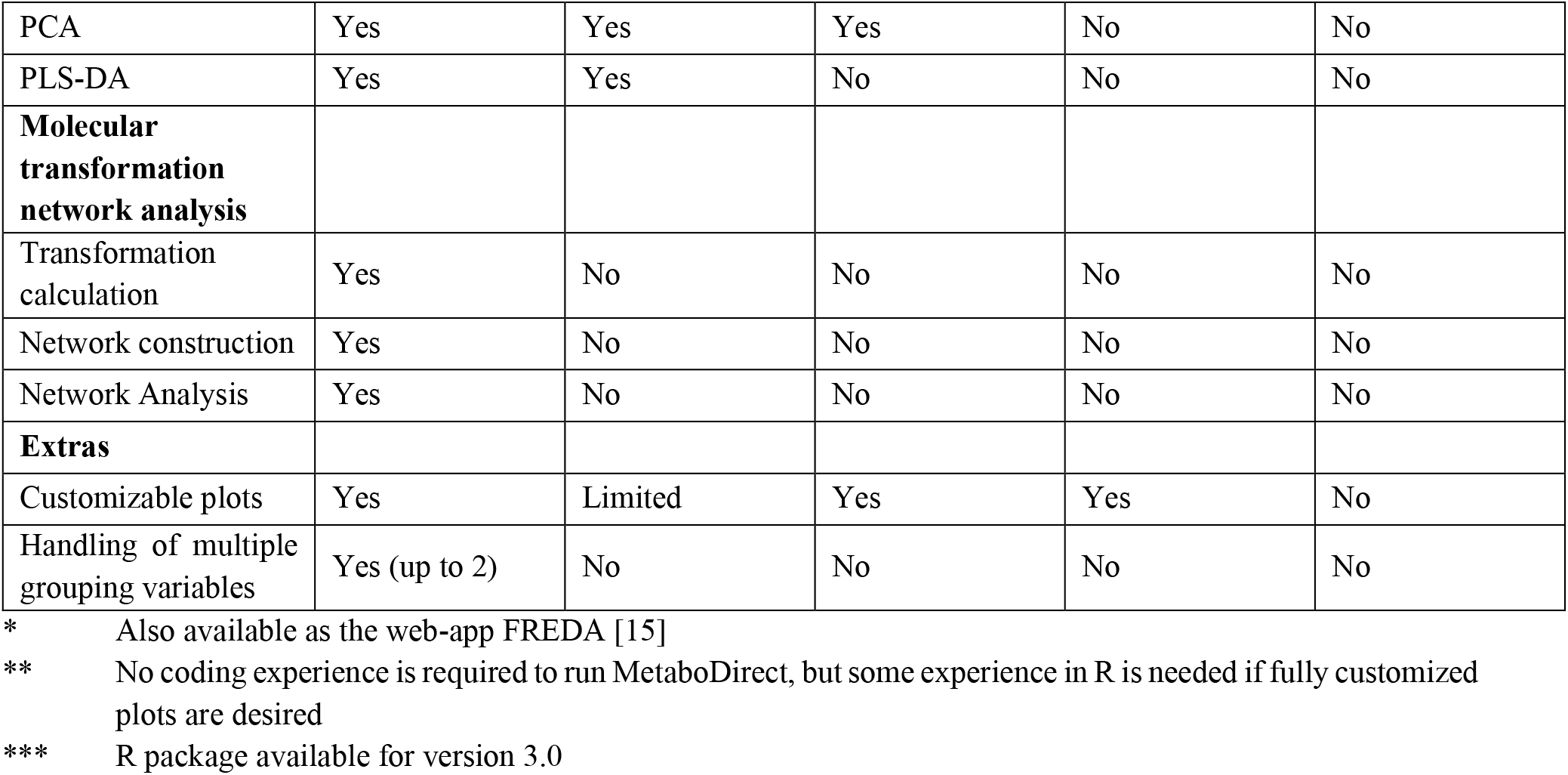
Comparison of the features of MetaboDirect with other available software for the analysis of FTICR-MS-based metabolomics datasets.

Here we introduce MetaboDirect, an easy-to-use, command-line based pipeline for the analysis of direct injection FTICR-MS-based metabolomics data collected from diverse ecosystems (soil, river, plants, bacterial cultures, etc.), which combines the easy usage of the web-based applications with the flexibility and customizability of the more programming-friendly software currently available. MetaboDirect accepts data produced by FTICR-MS (peak lists, peak intensities, metabolite molecular formula) or any other high resolution MS technique to facilitate data exploration, data visualization, chemodiversity, and statistical analysis. MetaboDirect is designed to run using a single line of code to automatically produce all the analyses, figures, and tables described in its documentation and its compute time is much faster than any other available FTICR-MS software. To further ease the access of MetaboDirect to scientists of all programming-skill levels, detailed information regarding it’s functioning, and available options is available through its User’s Guide (https://metabodirect.readthedocs.io/en/latest/).

In this manuscript, we showcase the use and outputs of MetaboDirect through the analysis of two FTICR-MS datasets. The first metabolomics dataset was generated from an established, ecologically relevant marine phage-host model system [34-36] designed to study new virus-host-nutrient interactions and its impact on the composition of bacterial metabolites [37]. The second metabolomics dataset was obtained from a study that aimed at elucidating plant leachate in particular *Sphagnum fallax* leachate degradation pathways and biochemical transformation in the presence and absence of microorganisms [12].

## IMPLEMENTATION

### 1. The MetaboDirect pipeline

The MetaboDirect pipeline is developed in Python 3.8 and R 4.0.2 [38] and is available to install through the Python Package Index (https://pypi.org/project/metabodirect/). It requires the Python dependencies NumPy [39], pandas [40, 41], seaborn [42], py4cytoscape and matplotlib [43]. The full documentation for the pipeline can be found on its ReadTheDocs webpage: https://metabodirect.readthedocs.io.

The MetaboDirect pipeline consists of 6 major steps/categories (Figure 1): (i) data pre-processing, (ii) data diagnostics, (iii) data exploration, (iv) chemodiversity analysis, (v) statistical analysis, and (vi) transformation network analysis. All these steps can be run directly with the “metabodirect” command. An additional script, “test_normalization” is included to help users select the best normalization method and can be run before the main MetaboDirect pipeline.

**Figure 1.**
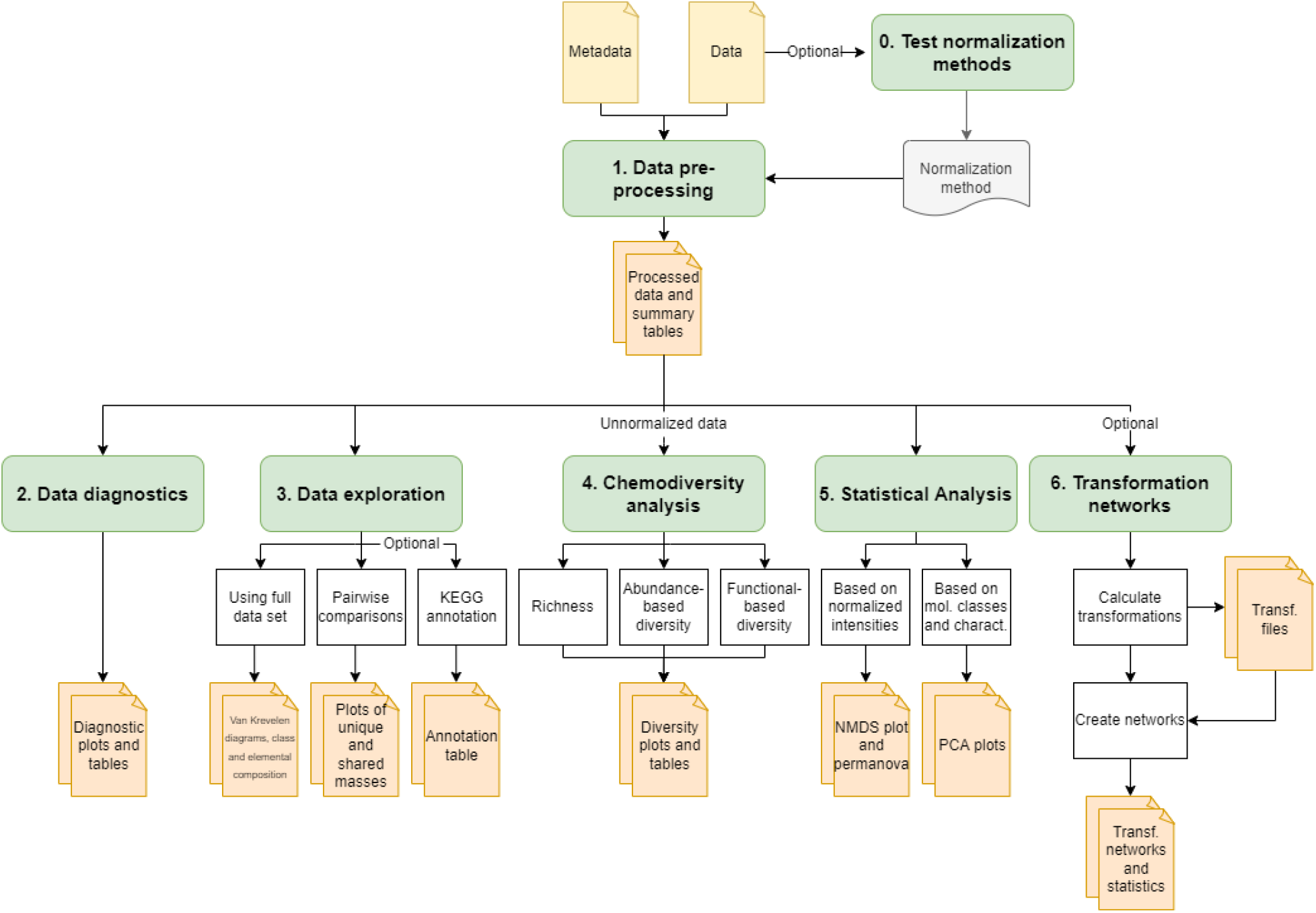
Workflow steps in the MetaboDirect pipeline and the outputs generated by each step.

MetaboDirect is designed to work directly with the .csv file generated by Formularity [24] from raw FTICR-MS data (.xml) which includes the list of assigned molecular formulas, their corresponding peak intensities (monoisotopic peaks) and *m/z* values. However, it can also be used with any list of masses and their corresponding molecular formulas as long as the .csv file is formatted according to the MetaboDirect documentation.

#### 1.0 Selection of the best normalization method

The companion script “test_normalization” uses the Statistical Procedure for the Analysis of Normalization Strategies (SPANS) [44] to aid in the selection of the best normalization method for the intensities of the detected peaks. This approach has been previously demonstrated to work well with FTICR-MS data [45] and uses a modified “spans_procedure” function from the R package *pmartR* [46]. The result of this step will be dependent on the nature of the dataset and the grouping variable that was analyzed. The calculated SPANS scores for the combination of normalization and subset methods are presented in a heatmap.

#### 1.1. Data pre-processing

This is the first step of the main MetaboDirect pipeline. During this step, detected peaks are filtered by their *m/z* values (based on the user’s input), isotopic presence (^13^C peaks), and error in formula assignment (0.5 ppm). Compound classes of each of the filtered peaks are then determined based on the assigned molecular formula using the criteria specified in Supplementary Table 1 [23, 47].

The molecular properties and hypothetical decomposability of the filtered peaks that received a molecular formula assignment are determined by calculating several thermodynamic and molecular indexes based on each peak’s elemental composition (equations used to calculate all thermodynamic indexes are included in Supplementary Table 2). These indexes include: nominal oxidation state of carbon (NOSC) that describes the average carbon oxidation state of the assigned peak based on its elemental composition [48]; Gibbs free energy (ΔG°C-ox or GFE) that indicates how likely the compound is to be degraded [48]; modified Aromaticity Index (AImod) that reflects the “density” of carbon-to-carbon double bonds within a molecule [49, 50]; and finally double bond equivalence (DBE) that represents the amount of unsaturation in a molecule and can indicate the presence of aromatic structures [50].

Furthermore, peak intensities are normalized in this step based on the user’s input. Normalization methods included in MetaboDirect are based on the normalization used in other similar tools [33, 45] and are detailed in Supplementary Table 3. This pre-processing step generates several .csv files containing the list of filtered peaks with their respective thermodynamic and molecular indexes and the normalized and unnormalized intensities which will be used in the next steps of the pipeline.

#### 1.2. Data diagnostics

This part of the pipeline generates plots of the total number of peaks detected in each sample after filtering (based on step 1.1) and the number of peaks that received a molecular formula assignment in each sample out of the total number of peaks. Both the total number of peaks and the total number of molecular formulas assigned per sample are reported in .csv tables and plotted as bar plots. Additionally, the data diagnostics step plots the error distribution of assigned molecular formulas as faceted scatterplots.

#### 1.3. Data exploration

This step produces several plots based on the molecular and thermodynamic properties of the peaks that received molecular formula assignment. During data exploration, MetaboDirect generates van Krevelen diagrams [51] of the peaks with molecular formulas; density and violin plots of the thermodynamic indices calculated in step 1.1, including whether or not there is a significant difference between the different groups using Tukey post hoc tests; bar plots with the molecular and elemental composition for each group/treatment; and finally pairwise comparison plots based on the user’s selected grouping variable/s. Pairwise comparisons are used to show which peaks are unique and which are shared between the different groups using van Krevelen diagrams and upset plots. As an additional option available in this step, MetaboDirect can use the assigned molecular formulas to query the KEGG database [52] using the R package *KEGGREST* [53] to provide putative KEGG Pathway, Module, and Brite annotations as .csv files.

#### 1.4. Chemodiversity analysis

For this step, raw peak intensities are sum-normalized and used to calculate commonly used chemodiversity metrics with functions from the R packages *vegan* [54] and *SYNCSA* [55]. Diversity metrics generated include species (metabolite) richness and rank abundance. Abundance-based diversity is measured with the Shannon diversity index [56], the Gini-Simpson index [57, 58], and the Chao1 richness estimator [59]. Functional based diversity, based on the compound’s elemental composition, potential decomposability, and insaturation/aromaticity, is measured with the Rao’s quadratic entropy index [60]. All diversity indexes are visualized as boxplots grouped by the user’s defined grouping variables and also exported as .csv files.

#### 1.5. Statistical analysis

In the data statistics step, the normalized intensities of the peaks (from step 1.1) are transformed into Bray-Curtis, Euclidean, or Jaccard distances (depending on the selected normalization method) using the “vegdist” function for the *vegan* package and then used to perform a permutational analysis of variance (PERMANOVA) [61]. Ordination of the data, based on the normalized intensities, is calculated using non-metric multidimensional scaling (NMDS) [62], as it provides a robust approach and can use any of the dissimilarity (distances) metrics mentioned before [63]. NMDS scores are then visualized as scatterplots using the first two components as axis, while the results of the PERMANOVA are exported as a .csv file. Additional ordination of the data is provided as Principal Component Analysis (PCA) [64] scree plots and biplots based on the molecular composition and magnitude-averaged thermodynamic indexes of the samples [65].

#### 1.6. Transformation networks

This is an optional step of the MetaboDirect pipeline as it is time consuming and, in most cases, is only needed to be run once per each dataset. Molecular transformation networks for each sample are generated in this step, where nodes represent peaks detected in the different samples and edges represent the putative chemical transformations happening between the nodes [21, 66, 67]. This step consists of two main processes: calculation of transformations and the creation of the molecular networks.

Transformations are determined by calculating the differences between the *m/z* of all peaks present in each sample and comparing them to a list of pre-defined masses (biochemical transformations key) [66] with a maximum error of 1 ppm. A user-defined mass list can also be used. The transformations included in the predefined biochemical transformation key are further classified in biotic or abiotic transformations [12]. These results are exported as .csv “edge” files containing the potential transformations occurring between the masses in each sample. Additional files with the number of transformations occurring per sample are also generated. Networks are then constructed using Cytoscape [68] and colored based on their molecular class. Transformation networks can then be used to inform the chemical connectives between the detected compounds. Furthermore, networks statistics will be calculated and reported as .csv tables and bar plots.

### 2. Pipeline testing

The performance of the MetaboDirect pipeline is tested here using two previously analyzed FTICR-MS datasets collected in negative ion-mode. The first came from the exametabolome of a marine phage-host model system that uses a known, ecologically relevant marine bacterium (*Pseudoalateromonas*) and two contrastingly different infecting phages (podovirus HP1 and siphovirus HS2) that have been extensively characterized via genomics and time-resolved transcriptomics and proteomics under nutrient-rich conditions [34-37]. Since viruses control microbes that provide essential ecosystem services through infection and reprogramming of the host cell metabolism, they can impact the composition of bacterial exometabolites into the ecosystem. This dataset is used here to show how MetaboDirect can help develop foundational approaches to studying how viral infection of bacterial communities could impact ecosystem outputs with significant impact on ecosystem functioning. Specifically, MetaboDirect was applied to determine how different infecting phages influence the metabolites produced under nutrient (P) rich conditions [37].

The second dataset was obtained from an incubation experiment of *S. fallax* leachate that was conducted in the presence and absence of microorganisms. This dataset was used to demonstrate how MetaboDirect can be used to help elucidate how organic matter degradation pathways can change in the presence or absence of microbial communities.

It is important to note that the goal of this paper is to provide two comprehensive examples of the application of the MetaboDirect pipeline and benchmark its “clock times” (i.e. compute time) against other currently available software, but not necessarily to provide biological interpretations of the data analyzed. Information regarding the incubation parameters, soil organic matter extraction protocols and high-resolution mass spectrometry data collection is provided in the Supplementary Information.

## RESULTS AND DISCUSSION

Here we demonstrate MetaboDirect’s functionality in data processing, filtering, and visualization (Figure 1) through the analysis of two distinct FT ICR-MS datasets. The bacterium-phage model system dataset included metabolites or mass spectrometry peaks present in 36 samples from bacteria infected by two different phages and a control treatment under nutrient rich conditions at different time points. The *S. fallax leachate* incubation metabolomics dataset consisted of 4 samples, two samples where the plant leachate was incubated anaerobically in the presence of in situ microbial communities and two samples where the plant leachate was incubated in the absence of any microbial communities.

### MetaboDirect pipeline testing

One of the advantages of the MetaboDirect pipeline is that it allows to automatically run all the analysis at once in a reproducible manner. The main steps of the MetaboDirect pipeline (steps I through V) run quickly, facilitating the rapid exploration of the data using different user-defined parameters in an efficient manner. When the compute time of MetaboDirect was benchmarked without its optional steps (KEGG mapping and the construction of transformation networks using datasets of different size), 40 samples were processed in less than one minute whereas 120 samples took as little as 2 mins to generate all the figures, plots and outputs discussed above at once. Even though the variation in execution time in MetaboDirect depends on both the number of samples analyzed, and the number of molecular formulas assigned per each sample, the MetaboDirect pipeline is still superior to other FTICR-MS software where the user is expected to plot the data one sample at time, and one figure type at a time.

For the bacterium-phage dataset a “report” file produced by Formularity [24] was processed through the whole MetaboDirect pipeline. The dataset had an average of 1025 peaks detected across the whole dataset (n = 36 samples) and an average of 495 peaks that got assigned a molecular formula. The main steps of the MetaboDirect pipeline (without KEGG database mapping or calculating transformation networks) took less than 1 minute (∼36 seconds) for this dataset. The analysis including KEGG mapping and calculation of the putative biochemical transformations took around 10 minutes. The full analysis including: KEGG mapping, calculating the putative biochemical transformations and creating the networks took about 21 minutes. For the *S. fallax* dataset, the data was obtained from the paper supplementary tables [12] and rearranged to fit the format needed for MetaboDirect. This was a smaller dataset (n = 4 samples) with an average of 1793 assigned molecular formulas across the whole dataset. For this dataset the main steps of the MetaboDirect pipeline were clocked at around 30 seconds. However, due to the larger number of assigned molecular formulas, full analysis of this dataset (including KEGG mapping and construction of molecular networks) took about 32 minutes.

Outside MetaboDirect, KEGG mapping is performed using KEGG MAPPER [69], a tool available directly through the KEGG website. While this tool searches various KEGG objects, including genes, KOs, EC numbers, metabolites, and drugs, against KEGG pathway maps, it requires the user to first identify the KEGG ID of each metabolite before manually importing the data to this MAPPER tool. MetaboDirect on the other hand performs all these steps automatically. The generation of biochemical transformations is currently done using the Cytoscape app MetaNetter_2 [70] that can only generate one network at a time and the user has to provide three files for each sample at a given time and manually set up network generation parameters such as mass tolerance. These files include: 1) a list of accurate masses per sample from the FTICR-MS data, 2) a list of accurate masses of common biochemical transformations that the user is interested in quantifying within the sample, 3) a metadata file that includes the different characteristics if each accurate mass. MetaboDirect on the other hand perform all these analysis at once and for all samples within a given dataset.

For both datasets, the first step was to use the “test_normalization” companion script to help determining which normalization method worked the best. FTICR data typically carry high biological or technical variation, and normalization is the first required step to enable data quantification. As proteomics analyses have shown [71] that normalization methods are dataset-dependent. MetaboDirect relies on the use of SPANS [44, 45] to identify the best normalization method for each dataset. For both datasets, median and zscore normalization, appeared to be the best normalization methods of this dataset as they had the highest SPANS score (Supplementary Figure 1). Thus, *median* normalization was selected for the bacterium-phage dataset and *zscore* for the *S. fallax* dataset for data normalization and statistical analysis in the subsequent steps.

Following normalization testing, the data pre-processing step was used to filter out specific peaks and to identify any problems within the dataset. In the bacterium-phage dataset, ∼200 masses were filtered out from the dataset because they were assigned molecular formulas containing an isotopic carbon. The diagnostics step further identified one sample with a very low number of assigned molecular formulas compared to the other samples, which can be a potential outlier for the analysis (Supplementary Figure 2). No peak was filtered in the *S. fallax* dataset it was previously filtered by the authors of that study.

Following the data pre-processing, the data exploration step was used to provide an overview of the molecular composition and thermodynamic characteristics of each of the user-defined groups for each dataset (the type of bacteriophage (HP1 vs. HS2 vs control), or the treatment (control vs inoculation)). The use of the molecular properties and thermodynamic indexes calculated by MetaboDirect are useful in studying how biotic and abiotic factors can influence the metabolites’ lability (NOSC and ΔG°C-ox) and degree of saturation of the metabolites present in each set of samples [20, 72].

Exploratory analysis of the bacterium-phage dataset showed that the chemical composition of the exometabolome of cells infected with the HS2 phage changed at 30 mins post inoculation after which it became more similar to the chemical composition of the uninfected cells (Supplementary Figure 3A), while the molecular composition of exametabolome from cells infected with the HP1 phage did not change throughout the experiment. This change in molecular composition resulted in changes in the modified aromaticity index (AI_mod) and the double bond equivalence (DBE), meaning that the number of aromatics compounds in HS2 exometabolome changed with time.

MetaboDirect automatically generates all pairwise comparisons for the selected grouping factors being analyzed. In this manner, the pipeline allows to identify which metabolites are common between the different conditions and which metabolites are unique to each treatment. For the bacterium-phage dataset, this analysis showed that most detected metabolites were shared between the infected and uninfected cells (Figure 2A) and that the unique metabolites were mostly protein-like, lignin-like, and lipid-like metabolites (Figure 2B and 2C). Conversely, pairwise comparisons of the *S. fallax* dataset showed that the number of unique metabolites in the control and the inoculated samples were almost the same as the number of shared metabolites (Supplementary Figure 4A), with lignin-like metabolites being most of the unique metabolites in the inoculated samples (Supplementary Figure 4B).

**Figure 2.**
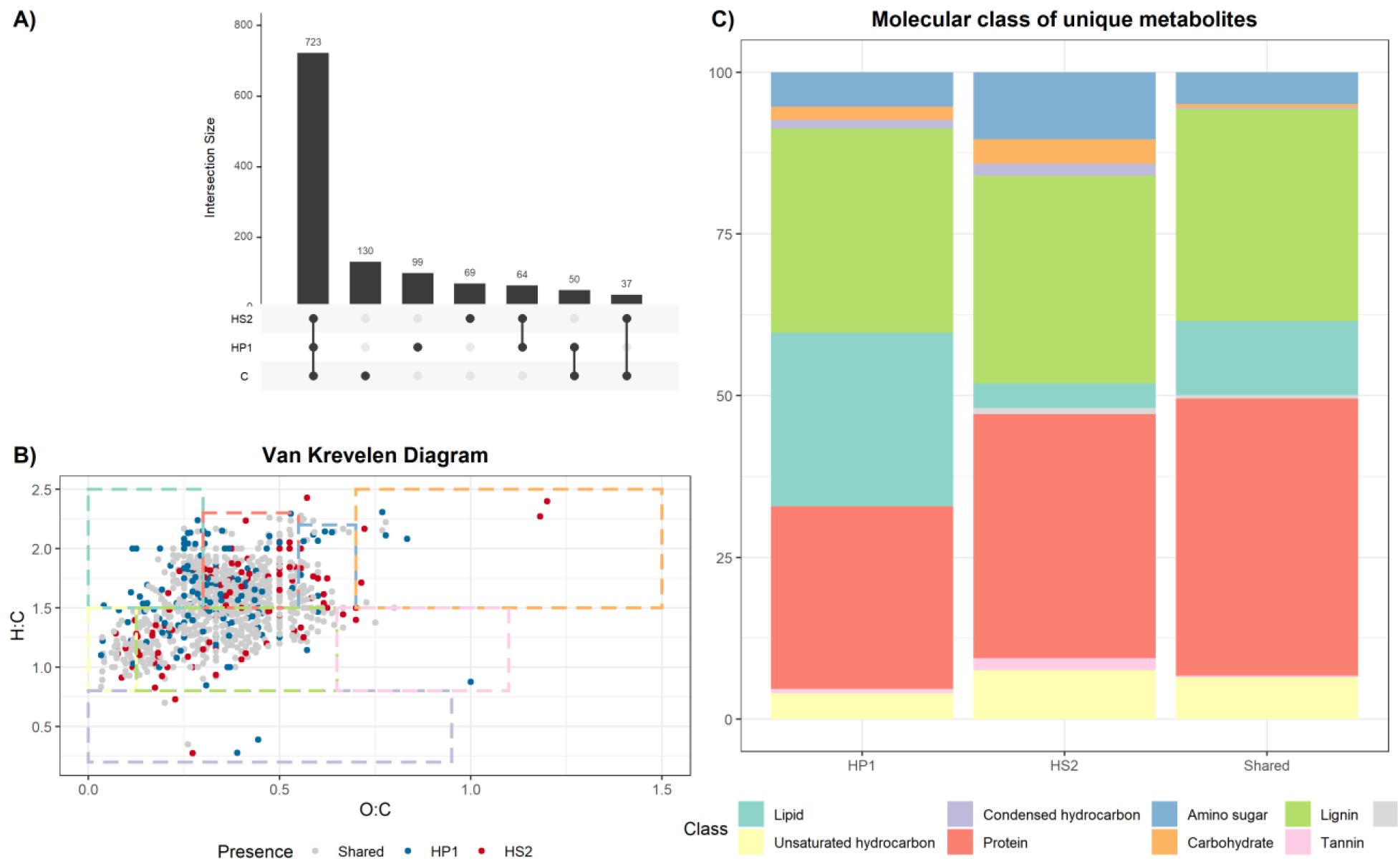
A) Upset plot showing the number of metabolites that are shared and unique between the uninfected and infected cells. B) Van Krevelen diagram showing metabolites that are shared and unique between cells infected with the two different phages, HP1 and HS2. C) Molecular composition of the unique metabolites showing that there are unique protein-like, lignin-like, and lipid-like metabolites in cells infected with each phage.

Diversity metrics, originally designed to study ecological species, can be adapted to analyze other systems, such as metabolites. In this case, molecular formulas are akin to individual species, while their peak intensity is used for abundance [73]. Commonly used diversity metrics in the study of metabolite assemblages include the measure of molecular richness (based only on the number of molecular formulas) [13, 14, 74]; the use of abundance-based diversity metrics such as Shannon, Gini-Simpson [73, 75, 76] or the Chao 1 [13, 14] indexes, which (based on the richness and the relative intensity of the metabolites); as well as the use Rao’s quadratic entropy to measure functional-based diversity [73, 76] (which uses different molecular characteristics as traits). MetaboDirect assists with calculating and visualizing all the diversity indexes mentioned above.

Chemodiversity analysis for the bacterium-phage dataset found little differences in the metabolite diversity between the infected and uninfected cells for either abundance-based diversity or functional diversity (Supplementary Figure 5A and 5C). For the *S. fallax* incubation dataset, inoculating the *S. fallax* leachate with microorganisms increased the diversity of the metabolites (i.e., richness) (Figure 5B) but decreased the functional diversity, suggesting that the metabolites in the inoculated samples were less diverse in terms of their decomposability or reactivity, aromaticity, and elemental composition (Figure 5D).

Multivariate analysis such as NMDS, PCA and PERMANOVA can be used to better understand the relationships of the normalized peak intensities or the molecular characteristics of the metabolites and the biotic and abiotic factors, as well as to find trends among large number of samples at the same time [13, 65, 77]. We used these methods to try to illustrate the effect of phage type and the time of the incubation on the relative amount and molecular characteristics of the metabolites produced by the bacterial-phage system during infection. The type of phage (or lack thereof) infecting the cells did not produce any significant differences in the metabolite/organic compound content of the samples (PERMANOVA p-value > 0.05, Supplementary 6C). The ordination analysis performed by MetaboDirect corroborated these results as it can be seen in the two ordination plots (Supplementary Figure6A and 6B).

The last step of MetaboDirect produces molecular transformation networks for each of the samples. These networks are generated *ab initio* from the masses that are determined through high-resolution mass spectrometry and are based on the fact that the ultra-high mass accuracy of the method allows identifying chemically transformed species using clearly defined mass differences [66, 67]. The number of transformations, converted to percentages can be used not only to quantify the differences in the microbial metabolic pathways between different groups, but can also assist in inferring the chemical identity of unknown metabolites (i.e. metabolites that didn’t receive a molecular formula) by identifying the connectivity between related metabolites since links in these networks correspond to actual chemical reactions and not metabolite correlation networks. The type of biochemical transformations on the other hand provides information on the type of reactions that are occurring within a given sample/system that take place either through enzymes whose presence can be validated through other omics analysis or through non-enzymatic metabolite interconversions within the cell/system that we usually do not account for in most microbiome studies. For the bacteriophage dataset, the most abundant chemical transformation was methylation (i.e. loss or gain of a methyl group). However, we didn’t observe significant changes in the quality of quantity of all biochemical transformations with infection. Interestingly, analysis of the transformation networks produced by MetaboDirect for the bacterium-phage dataset showed that for most of the samples, there was always a cluster of protein-like metabolites interacting with phenol (lignin)-like metabolites (Figure 3B) suggesting that these *ab initio* networks will be very useful for future scientific discoveries especially when such studies are complemented with other omics datasets.

**Figure 3.**
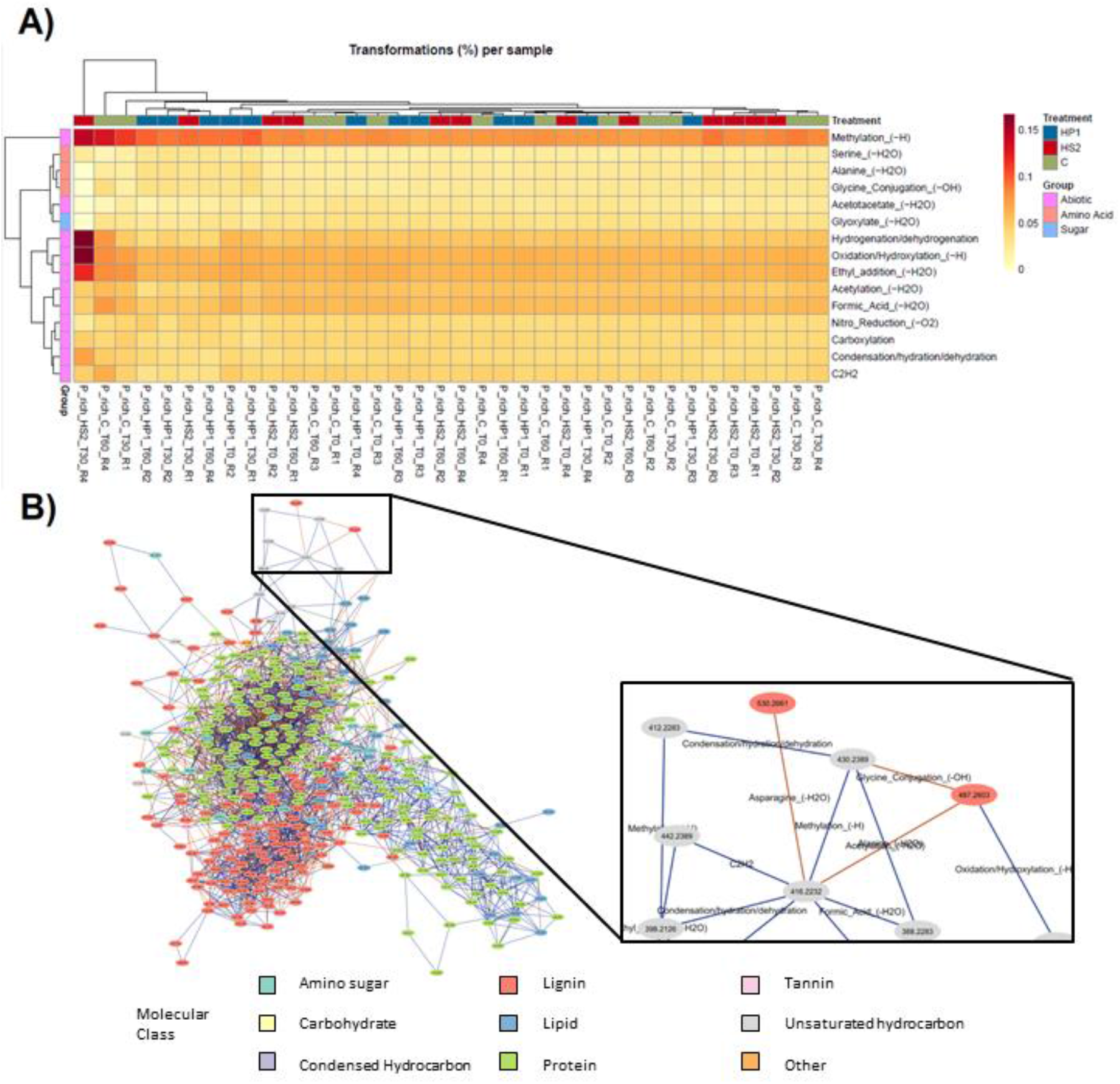
A) Heatmap with the top 15 more abundant transformations among all the samples converted into percentage of the total number of transformations in each sample. B) Molecular network for the sample P_rich_C_T30_R3. Like most of the other samples, the transformation network showed a cluster of lignin-like metabolites interacting with a cluster of protein-like metabolites. The zoomed panel at the left shows that the transformation networks have the masses of each sample as nodes, while the edges between those nodes represent the transformations. Transformations are classified into biotic (sugar and amino acid transformations) or abiotic transformations as discussed in [9].

### Comparison with other available software

MetaboDirect was designed for reproducible analysis of direct injection FTICR-MS data, ranging from diagnostics and data exploration to chemodiversity and statistical analysis (Table 1), and it is fully automated with a single line of code and offers an automated pipeline for the generation and visualization of transformation networks. As discussed above, this type of analysis is currently performed through the use of the Cytoscape plugin MetaNetter_2 [70], in a laborious and time-consuming process which involves exhaustive file preparation as it must be done on a per sample basis. While some software packages with a GUI, such as the web-based apps MetaboAnalyst, UltraMassExplorer and FREDA are user-friendly, they are restricted to some of the most common analysis tools for FTICR-MS data such as the generation of van Krevelen plots and multivariate statistical analysis, and they lack customization. On the other hand, more comprehensive software packages, which are presented in the form of libraries for commonly used programming languages, such as *PyKrev* and *ftmsRanalysis*, allow the users to deeply customize their data analysis. However, they require medium to advanced skills in computer programming to be able to take full advantage of their parametrization.

MetaboDirect stands in the middle of those groups, requiring minimal coding experience, as all the output files are obtained using a single line of code. Moreover, MetaboDirect provides the user with all the R scripts used in the generation of all the tables and visualizations, thus allowing the user to fully customize any of the figures and tables produced by MetaboDirect and to use that data in any additional analysis. Therefore, MetaboDirect can be attractive to users with all programming skill levels, allowing them to take advantage of a fully automated pipeline that can be easily customizable if needed.

As observed in Table 1, MetaboDirect can perform all the analysis offered by the other available software for FTICR-MS data, except for “Raw spectra processing” and “Molecular formula assignment”. Nonetheless, these first two steps in the processing of FTICR-MS data can be easily achieved by using Formularity or CoreMS.

## CONCLUSION

The use of high-resolution mass spectrometry, specifically FTICR-MS, to characterize the molecular composition of NOM in different systems is increasing quickly, and thus the development of reproducible open-source tools for the analysis of such data is urgently needed. Here we present MetaboDirect, a user-friendly, accessible, and highly comprehensive tool for scientists working to characterize how different biotic and abiotic factors influence organic compound composition in diverse settings and systems that can be used to provide a quick overview of the data, upon which more in-depth analysis can be built. The highly reproducible nature of the analysis provided by MetaboDirect, coupled with the detailed user manual, will allow scientists of all skill levels to fully explore and work with FTICR-MS data. This in return will greatly facilitate the integration of metabolomics within current microbiome studies and advance our knowledge of how microbial communities influence and are influenced by the chemical makeup of the surrounding system.

## AVAILABILITY AND REQUIREMENTS

**Project name:** MetaboDirect

**Project home page:** https://github.com/Coayala/MetaboDirect

**Operating system(s):** Windows, iOS, Linux

**Programming language:** Python, R

**Other requirements:** Cytoscape 3.8 or higher

**License:** MIT License

### LIST OF ABBREVIATIONS

DOM: Dissolved organic matter
FTICR-MS: Fourier-transform ion cyclotron resonance mass spectrometry
GUI: Graphical user interface
NMDS: Non-metric multidimensional scaling
NOM: Natural organic matter
PCA: Principal component analysis
SOM: Soil organic matter

## DECLARATIONS

### Ethics approval and consent to participate

Not applicable.

### Consent for publication

Not applicable.

### Availability of data and materials

(1) The bacterium – phage system dataset is available here https://doi.org/10.17605/OSF.IO/XFHZ9; (2) The original *Sphagnum* dataset is available here [12] (https://agupubs.onlinelibrary.wiley.com/action/downloadSupplement?doi=10.1029%2F2020JG006079&file=2020JG006079-sup-0007-Table+SI-S02.xlsx) and the formatted one according to MetaboDirect format is available here https://doi.org/10.17605/OSF.IO/XFHZ9

The code of the MetaboDirect pipeline used for this analysis is available in its GitHub repository (https://github.com/Coayala/MetaboDirect) with DOI: https://doi.org/10.5281/zenodo.5874805“ https://doi.org/10.5281/zenodo.5874805.

### Author’s contributions

MT, NGG, YC and NC designed the study; MT, NGG, YC, NC and MS obtained the funding; JF, CHV and MD performed the experiments and processed samples; CAO analyzed the data used for the manuscript; CAO and NGG developed the software used for the analysis; VFZ, GH, RA tested and validated the use of the software; CAO and MT wrote the manuscript. All authors contributed to manuscript review and editing.

## Acknowledgements

We thank members of the Tfaily Lab, Sullivan Lab and EMSL staff for feedback on design and implementation of MetaboDirect, as well as comments on the manuscript.

## Author’s information

NGG: Current position – Roche, Pleasanton, CA, 94588, USA. Current JF: Current position – University of California, Davis | Department of Plant Pathology, Davis, CA 95616-8751. Current

## REFERENCES

1. Cavicchioli, R., et al., Scientists’ warning to humanity: microorganisms and climate change. Nature Reviews Microbiology, 2019. 17(9): p. 569–586.

2. Mohanty, B., et al., Biogeochemical Cycles in Soil Microbiomes in Response to Climate Change, in Climate Change and the Microbiome: Sustenance of the Ecosphere, D.K. Choudhary, A. Mishra, and A. Varma, Editors. 2021, Springer International Publishing: Cham. p. 491–519.

3. Malik, A. and G. Gleixner, Importance of microbial soil organic matter processing in dissolved organic carbon production. FEMS Microbiology Ecology, 2013. 86(1): p. 139–148.

4. Myneni, S.C., Chemistry of natural organic matter—The next step: Commentary on a humic substances debate. Journal of environmental quality, 2019. 48(2): p. 233–235.

5. Wu, X., et al., Microbial Interactions With Dissolved Organic Matter Drive Carbon Dynamics and Community Succession. Frontiers in Microbiology, 2018. 9(1234).

6. McDowell, W.H., Dissolved organic matter in soils—future directions and unanswered questions. Geoderma, 2003. 113(3): p. 179–186.

7. Castro, H.F., et al., Soil microbial community responses to multiple experimental climate change drivers. Applied and environmental microbiology, 2010. 76(4): p. 999–1007.

8. Soule, M.C.K., et al., Environmental metabolomics: Analytical strategies. Marine Chemistry, 2015. 177: p. 374–387.

9. Kujawinski, E.B., The Impact of Microbial Metabolism on Marine Dissolved Organic Matter. Annual Review of Marine Science, 2011. 3(1): p. 567–599.

10. AminiTabrizi, R., et al., Elevated temperatures drive abiotic and biotic degradation of organic matter in a peat bog under oxic conditions. Science of The Total Environment, 2021: p. 150045.

11. Fudyma, J.D., et al., Coupled biotic-abiotic processes control biogeochemical cycling of dissolved organic matter in the Columbia River hyporheic zone. Frontiers in Water, 2021. 2(PNNL-SA-158822).

12. Fudyma, J.D., et al., Sequential Abiotic-Biotic Processes Drive Organic Carbon Transformation in Peat Bogs. Journal of Geophysical Research: Biogeosciences, 2021. 126(2): p. e2020JG006079.

13. Kellerman, A.M., et al., Chemodiversity of dissolved organic matter in lakes driven by climate and hydrology. Nature Communications, 2014. 5(1).

14. Li, X.-M., et al., Molecular Chemodiversity of Dissolved Organic Matter in Paddy Soils. Environmental Science & Technology, 2018. 52(3): p. 963–971.

15. Villas-Bôas, S.G., et al., Mass spectrometry in metabolome analysis. Mass spectrometry reviews, 2005. 24(5): p. 613–646.

16. Hertkorn, N., et al., Natural Organic Matter and the Event Horizon of Mass Spectrometry. Analytical Chemistry, 2008. 80(23): p. 8908–8919.

17. Sleighter, R.L. and P.G. Hatcher, The application of electrospray ionization coupled to ultrahigh resolution mass spectrometry for the molecular characterization of natural organic matter. Journal of Mass Spectrometry, 2007. 42(5): p. 559–574.

18. Cooper, W.T., et al., A history of molecular level analysis of natural organic matter by FTICR mass spectrometry and the paradigm shift in organic geochemistry. Mass Spectrometry Reviews, 2020.

19. González-Domínguez, R., A. Sayago, and Á. Fernández-Recamales, Direct infusion mass spectrometry for metabolomic phenotyping of diseases. Bioanalysis, 2017. 9(1): p. 131–148.

20. AminiTabrizi, R., et al., Controls on soil organic matter degradation and subsequent greenhouse gas emissions across a permafrost thaw gradient in Northern Sweden. Frontiers in Earth Science, 2020. 8(PNNL-SA-156978).

21. Fudyma, J.D., et al., Untargeted metabolomic profiling of Sphagnum fallax reveals novel antimicrobial metabolites. Plant Direct, 2019. 3(11): p. e00179.

22. Mann, B.F., et al., Indexing Permafrost Soil Organic Matter Degradation Using High-Resolution Mass Spectrometry. PLOS ONE, 2015. 10(6): p. e0130557.

23. Tfaily, M.M., et al., Advanced Solvent Based Methods for Molecular Characterization of Soil Organic Matter by High-Resolution Mass Spectrometry. Analytical Chemistry, 2015. 87(10): p. 5206–5215.

24. Tolić, N., et al., Formularity: Software for Automated Formula Assignment of Natural and Other Organic Matter from Ultrahigh-Resolution Mass Spectra. Analytical Chemistry, 2017. 89(23): p. 12659–12665.

25. Corilo, Y., W. Kew, and L.A. McCue, EMSL-Computing/CoreMS: CoreMS 1.0.0 (v1.0.0). 2021.

26. Bramer, L.M., et al., ftmsRanalysis: An R package for exploratory data analysis and interactive visualization of FT-MS data. PLOS Computational Biology, 2020. 16(3): p. e1007654.

27. Leefmann, T., S. Frickenhaus, and B.P. Koch, UltraMassExplorer: a browser-based application for the evaluation of high-resolution mass spectrometric data. Rapid Communications in Mass Spectrometry, 2019. 33(2): p. 193–202.

28. Pacific Northwest National Laboratory. FREDA. 2021 [cited 2021 August 6]; Available from: https://msc-viz.emsl.pnnl.gov/FREDA/.

29. Pang, Z., et al., MetaboAnalyst 5.0: narrowing the gap between raw spectra and functional insights. Nucleic Acids Research, 2021. 49(W1): p. W388–W396.

30. Rosa, T.R., et al., DropMS: Petroleomics Data Treatment Based in Web Server for High-Resolution Mass Spectrometry. Journal of the American Society for Mass Spectrometry, 2020. 31(7): p. 1483–1490.

31. Kew, W., et al., Interactive van Krevelen diagrams – Advanced visualisation of mass spectrometry data of complex mixtures. Rapid Communications in Mass Spectrometry, 2017. 31(7): p. 658–662.

32. Brockman, S.A., E.V. Roden, and A.D. Hegeman, Van Krevelen diagram visualization of high resolution-mass spectrometry metabolomics data with OpenVanKrevelen. Metabolomics, 2018. 14(4): p. 48.

33. Kitson, E., et al., PyKrev: A Python Library for the Analysis of Complex Mixture FT-MS Data. Journal of the American Society for Mass Spectrometry, 2021. 32(5): p. 1263–1267.

34. Duhaime, M.B., et al., Comparative Omics and Trait Analyses of Marine Pseudoalteromonas Phages Advance the Phage OTU Concept. Frontiers in Microbiology, 2017. 8(1241).

35. Duhaime, M.B., A. Wichels, and M.B. Sullivan, Six <i>Pseudoalteromonas</i> Strains Isolated from Surface Waters of Kabeltonne, Offshore Helgoland, North Sea. Genome Announcements, 2016. 4(1): p. e01697–15.

36. Duhaime, M.B., et al., Ecogenomics and genome landscapes of marine Pseudoalteromonas phage H105/1. The ISME journal, 2011. 5(1): p. 107–121.

37. Howard-Varona, C., et al., Phage-specific metabolic reprogramming of virocells. The ISME Journal, 2020. 14(4): p. 881–895.

38. R Core Team, R: A Language and Environment for Statistical Computing. 2020.

39. Harris, C.R., et al., Array programming with NumPy. Nature, 2020. 585: p. 357–362.

40. McKinney, W. Data structures for statistical computing in python. 2010.

41. The pandas development team, pandas-dev/pandas: Pandas. 2020.

42. Michael, L.W., seaborn: statistical data visualization. Journal of Open Source Software, 2021. 6(60): p. 3021.

43. Hunter, J.D., Matplotlib: A 2D graphics environment. Computing in Science \& Engineering, 2007. 9: p. 90--95.

44. Webb-Robertson, B.J.M., et al., A statistical selection strategy for normalization procedures in LC-MS proteomics experiments through dataset-dependent ranking of normalization scaling factors. Proteomics, 2011. 11(24): p. 4736–4741.

45. Thompson, A.M., et al., Fourier transform ion cyclotron resonance mass spectrometry (FT-ICR-MS) peak intensity normalization for complex mixture analyses. Rapid Communications in Mass Spectrometry, 2021. 35(9): p. e9068.

46. Stratton, K.G., et al., pmartR: Quality Control and Statistics for Mass Spectrometry-Based Biological Data. Journal of Proteome Research, 2019. 18(3): p. 1418–1425.

47. Tfaily, M.M., et al., Elevated [CO2] changes soil organic matter composition and substrate diversity in an arid ecosystem. Geoderma, 2018. 330: p. 1–8.

48. LaRowe, D.E. and P. Van Cappellen, Degradation of natural organic matter: A thermodynamic analysis. Geochimica et Cosmochimica Acta, 2011. 75(8): p. 2030–2042.

49. Chassé, A.W., et al., Chemical force spectroscopy evidence supporting the layer-by-layer model of organic matter binding to iron (oxy) hydroxide mineral surfaces. Environmental science & technology, 2015. 49(16): p. 9733–9741.

50. Koch, B.P. and T. Dittmar, From mass to structure: An aromaticity index for high-resolution mass data of natural organic matter. Rapid communications in mass spectrometry, 2006. 20(5): p. 926–932.

51. Kim, S., R.W. Kramer, and P.G. Hatcher, Graphical method for analysis of ultrahigh-resolution broadband mass spectra of natural organic matter, the van Krevelen diagram. Analytical chemistry, 2003. 75(20): p. 5336–5344.

52. Kanehisa, M. and S. Goto, KEGG: Kyoto Encyclopedia of Genes and Genomes. Nucleic Acids Research, 2000. 28(1): p. 27–30.

53. Tenenbaum, D. and B.P. Maintainer, KEGGREST: Client-side REST access to the Kyoto Encyclopedia of Genes a nd Genomes (KEGG).

54. Oksanen, J., et al., vegan: Community Ecology Package.

55. Debastiani, V. and V. Pillar, SYNCSA - R tool for analysis of metacommunities based on functional traits and phylogeny of the community components. Bioinformatics, 2012. 28: p. 2067–2068.

56. Shannon, C.E., A mathematical theory of communication. The Bell System Technical Journal, 1948. 27(3): p. 379–423.

57. Gini, C., Variabilità e mutabilità. Reprinted in Memorie di metodologica statistica (Ed. Pizetti E, 1912.

58. Simpson, E.H., Measurement of diversity. nature, 1949. 163(4148): p. 688–688.

59. Chao, A. and T.-J. Shen, Nonparametric estimation of Shannon’s index of diversity when there are unseen species in sample. Environmental and ecological statistics, 2003. 10(4): p. 429–443.

60. Rao, C.R., Diversity and dissimilarity coefficients: a unified approach. Theoretical population biology, 1982. 21(1): p. 24–43.

61. Anderson, M.J., Permutational Multivariate Analysis of Variance (PERMANOVA). Wiley StatsRef: Statistics Reference Online, 2017: p. 1–15.

62. Kruskal, J.B., Nonmetric multidimensional scaling: a numerical method. Psychometrika, 1964. 29(2): p. 115–129.

63. Holland, S.M., Non-metric multidimensional scaling (MDS). Department of Geology, University of Georgia, Athens, Tech. Rep. GA, 2008: p. 30602–2501.

64. Wold, S., K. Esbensen, and P. Geladi, Principal component analysis. Chemometrics and intelligent laboratory systems, 1987. 2(1-3): p. 37–52.

65. Miao, Y., et al., Molecular characterization of root exudates using Fourier transform ion cyclotron resonance mass spectrometry. Journal of Environmental Sciences, 2020. 98: p. 22–30.

66. Breitling, R., et al., Ab initio prediction of metabolic networks using Fourier transform mass spectrometry data. Metabolomics, 2006. 2(3): p. 155–164.

67. Longnecker, K. and E.B. Kujawinski, Using network analysis to discern compositional patterns in ultrahigh-resolution mass spectrometry data of dissolved organic matter. Rapid Communications in Mass Spectrometry, 2016. 30(22): p. 2388–2394.

68. Shannon, P., et al., Cytoscape: a software environment for integrated models of biomolecular interaction networks. Genome research, 2003. 13(11): p. 2498–2504.

69. Cantu, D.C., Y. Chen, and P.J. Reilly, Thioesterases: A new perspective based on their primary and tertiary structures. Protein Science, 2010. 19(7): p. 1281–1295.

70. Jourdan, F., et al., MetaNetter: inference and visualization of high-resolution metabolomic networks. Bioinformatics, 2007. 24(1): p. 143–145.

71. Callister, S.J., et al., Normalization approaches for removing systematic biases associated with mass spectrometry and label-free proteomics. Journal of proteome research, 2006. 5(2): p. 277–286.

72. Lv, J., et al., Molecular-scale investigation with ESI-FT-ICR-MS on fractionation of dissolved organic matter induced by adsorption on iron oxyhydroxides. Environmental science & technology, 2016. 50(5): p. 2328–2336.

73. Mentges, A., et al., Functional Molecular Diversity of Marine Dissolved Organic Matter Is Reduced during Degradation. Frontiers in Marine Science, 2017. 4(194).

74. Ding, Y., et al., Chemodiversity of soil dissolved organic matter. Environmental science & technology, 2020. 54(10): p. 6174–6184.

75. Seidel, M., et al., Molecular-level changes of dissolved organic matter along the Amazon River-to-ocean continuum. Marine Chemistry, 2015. 177: p. 218–231.

76. Tanentzap, A.J., et al., Chemical and microbial diversity covary in fresh water to influence ecosystem functioning. Proceedings of the National Academy of Sciences, 2019. 116(49): p. 24689–24695.

77. Sleighter, R.L., et al., Multivariate statistical approaches for the characterization of dissolved organic matter analyzed by ultrahigh resolution mass spectrometry. Environmental science & technology, 2010. 44(19): p. 7576–7582.

