## Supplementary Information for "MetaboDirect: An Analytical Pipeline for the processing of FTICR-MS-based Metabolomics Data"

[Supplementary Figure 3. A) Changes in the molecular class composition of the bacterium-phage exometabolome during the incubation. A reduction in the percentage of lignins is observed at 30 minutes after inoculation. B) Violin plot of the changes in the aromatic index reflect the changes in molecular composition, cells infected with the HS2 phage have lower AImod at 30 minutes after inoculation (Tukey HSD test, p-value < 0.05). C) Violin plot showing that double bond equivalence (DBE) of HS2 is reduced after 30 minutes and remains low until the end of the experiment (Tukey HSD test, p-value < 0.05). For B), C) and D) * (p-value < 0.05), ** (p-value < 0.01), *** (p-value < 0.001), **** (p-value < 0.0001).	12](#_Toc93664481)

### SUPPLEMENTARY METHODS

**Bacterium-phage data set**

The experimental setup from which the data was obtained consisted of a total of three treatments with twelve replicates each (4 replicates x 3 time points). Treatments included bacteria treated with one of two bacteriophages (HP1 or HS2) or with neither of them (control) growing under nutrient (P) rich conditions. Growth and infection were conducted as described previously [31, 67]. Briefly, *Pseudoalteromonas sp* 13-15 were grown shaking at 150 rpm at 21°C in wither 1%Z+CNP media for P-rich or 1%Z+CN for P-poor. These consisted of 26g sea salts per L, 1% Zobell (26g sea salts, 1g yeast extract, 5g proteose peptone per L), 8.3mM ammonium sulfate, and 0.15mM phosphoric acid (in P-rich medium only), with 11mM glucose added after autoclaving. One colony was inoculated into 10mL and grown overnight before 5x10^8^ cells were transferred to 200mL in 1L flasks and grown to mid- to late-exponential phase. Then, 1x10^8^ cells were transferred in triplicate to a 1.5mL tube and the volume was adjusted to 1mL. samples were incubated for 15 min after phage addition at a multiplicity of infection of ~5, and infections were diluted 10-fold in a 1L bottle with fresh medium. From the infections and controls, three samples were collected at 0-, 30- and 60-min post dilution of infection and were 0.2-uM filtered.

**Sample preparation.** Filtered exometabolome aqueous samples were subjected to solid phase extraction (SPE) to remove any salts that can interfere with ionization during mass spectrometry analysis. Briefly, samples were acidified to pH 2 using 1M HCl to enhance extraction efficiency. Acidified samples were then passed through a 3 mL Bond Elut PPE cartridge (Agilent), previously prepped with 3 mL of laboratory grade MeOH, connected to a vacuum. After passing all the samples, cartridges were washed with 3 mL of a 0.01 M HCl solution. The washing step was repeated five times. Washed cartridges were removed from the vacuum and dried using filtered air. Finally, samples were eluted into 2 mL glass vials with 1.5 mL of MeOH and stored at -80 °C until used.

**Direct injection FT-ICR-MS Data Acquisition.** A Bruker 9.4-Tesla, coupled to a standard Bruker electrospray ionization (ESI) source, was used to collect high-resolution mass spectrometry data of the exometabolome filtrates. Before data collection, the instrument was tuned using a Suwannee River fulvic acid (SRFA) standard, purchased from the International Humic Substances Society (IHCC). All samples, standards, and blanks (HPLC grade methanol) were injected directly to the ESI source. A flushing with a mixture of water and methanol was performed in between each sample to prevent carry over. Ion accumulation time (IAT) varied from sample to sample to account for differences in C content. Spectra for each sample was obtained as an average of 144 individual scans. DataAnalysis, version 4.2, (BrukerDaltonik) was used to extract a list of *m/z* values from raw spectra files using the FTICR-MS peak picker module with a signal-to-noise ratio (S/N) threshold of 7 and the default absolute intensity threshold of 100. Spectra were internally calibrated using an organic matter homologous series separated by 14 Da (CH_2_ groups).

**Formula assignment.** Molecular formula assignment was done using the software package Formularity [12]. Formulas were assigned using the following criteria: S/N > 7, mass measurement error < 1 ppm, C, H, O, N, P and S as the only viable elements, and P required the presence of at least four O. For peaks with large mass ratios (*m/z* > 500 Da), formulas were assigned through propagation of CH_2_, O and H_2_ homologous series. For peaks with multiple molecular formula candidates, the formula with the lowest error and the lowest number of heteroatoms was picked.

### SUPPLEMENTARY TABLES

#### Supplementary Table 1. O/C and H/C ratios used to assign putative molecular classes to the detected metabolites

| **Molecular Class** | **O/C ratio** | | **H/C ratio** | |
| --- | --- | --- | --- | --- |
|  | **Min** | **Max** | **Min** | **Max** |
| Lipid | 0 | 0.3 | 1.5 | 2.5 |
| Unsaturated hydrocarbon | 0 | 0.125 | 0.8 | 2.5 |
| Protein | 0.3 | 0.55 | 1.5 | 2.3 |
| Amino sugar | 0.55 | 0.7 | 1.5 | 2.2 |
| Carbohydrate | 0.7 | 1.5 | 1.5 | 2.5 |
| Lignin | 0.125 | 0.65 | 0.8 | 1.5 |
| Tannin | 0.65 | 1.1 | 0.8 | 1.5 |
| Condensed hydrocarbon | 0 | 0.95 | 0.2 | 0.8 |

#### Supplementary Table 2**.** Equations used to calculate thermodynamic and molecular indexes based on the assigned molecular of the detected mass spectrometry peaks based on their *m/z* values

| **Thermodynamic Index** | **Formula** |
| --- | --- |
| Nominal oxidation state of carbon (NOSC) | $NOSC= -\frac{4C+H-3N-2O+5P-2S}{C}+4$ |
| Gibbs free energy (ΔG°C-ox) | $\Delta G^{\circ}C=60.3-28.5*NOSC$ |
| Modified aromaticity index (AImod) | $AImod=1+C-\frac{O}{2}-S-\frac{\frac{H+P+N}{2}}{C-O-S-N-P}$ |
| Double bond equivalence (DBE) | $DBE=1+\frac{2C-H+P+N}{2}$ |

#### Supplementary Table 3. Normalization methods available in MetaboDirect

| **Normalization Method** | **Formula** |
| --- | --- |
| max | ${NormIntensity}_{i,j}=\frac{I_{i, j}}{{max(I)}_{i}}$ |
| minmax | ${NormIntensity}_{i,j}=\frac{I_{i, j}-{min(I)}_{i}}{{max(I)}_{i}-{min(I)}_{i}}$ |
| mean | ${NormIntensity}_{i,j}=\frac{I_{i, j}-{mean(I)}_{i}}{{max(I)}_{i}-{min(I)}_{i}}$ |
| median | ${NormIntensity}_{i,j}=\frac{I_{i, j}-{median(I)}_{i}}{{max(I)}_{i}-{min(I)}_{i}}$ |
| sum | ${NormIntensity}_{i,j}=\frac{I_{i, j}}{\sum I_{i}}$ |
| zscore | ${NormIntensity}_{i,j}=\frac{I_{i, j}-{mean(I)}_{i}}{{std.dev(I)}_{i}}$ |
| binary | $presence=1; absence=0$ |

### SUPPLEMENTARY FIGURES

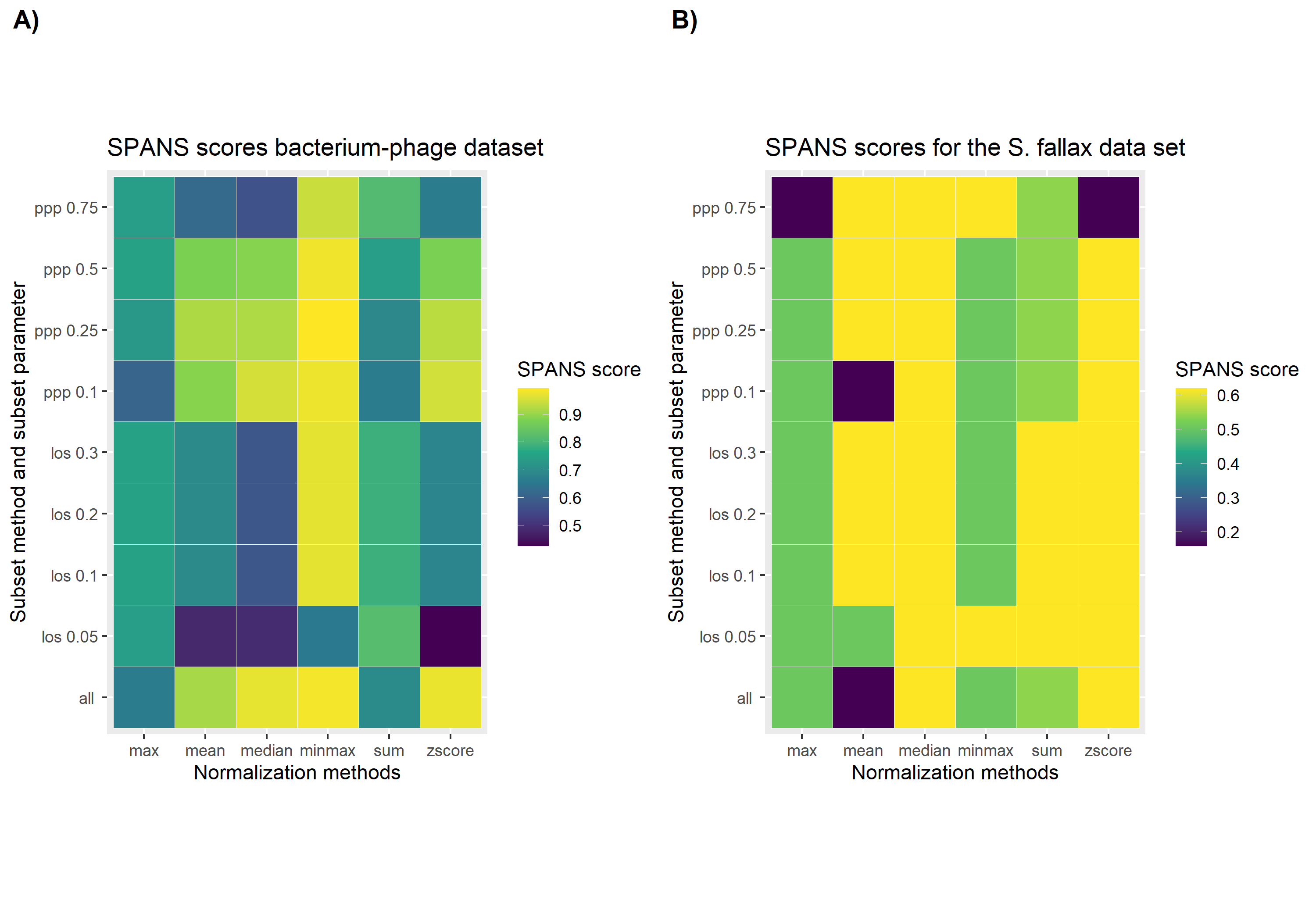

#### Supplementary Figure 1. SPANS score calculated by the “test_normalization” companion script for A) the bacterium-phage data set and B) the *Sphagnum fallax* data set. The x-axis shows the available normalization methods within MetaeboDirect, while the y-axis shows multiple combinations of the subset methods with different subset parameters. The SPANS score is shownshowed as a color scale with yellow as the highest score. For more information consult the User’s Guide (https://metabodirect.readthedocs.io).

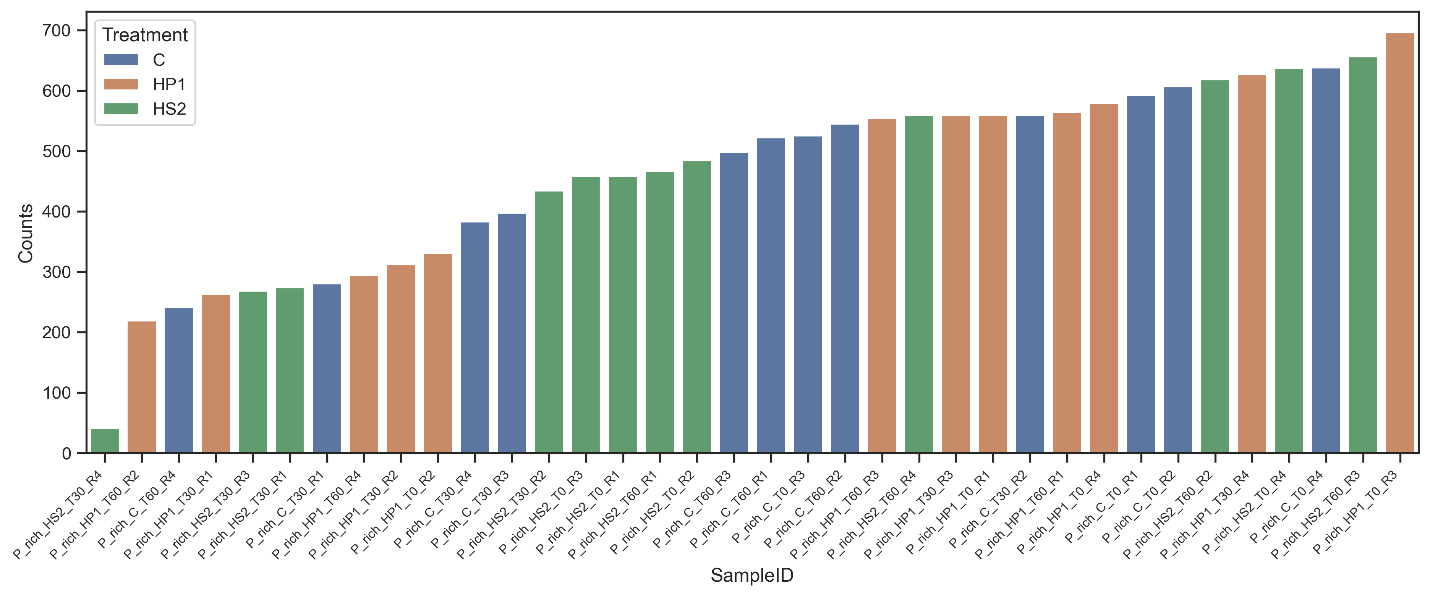

#### Supplementary Figure 2. Number of detected peaks that were assigned a molecular formula in the bacterium-phage dataset. The sample P_rich_HS2_T30_R4 has few peaks that were assigned a molecular formula and can be a potential outlier during the study.

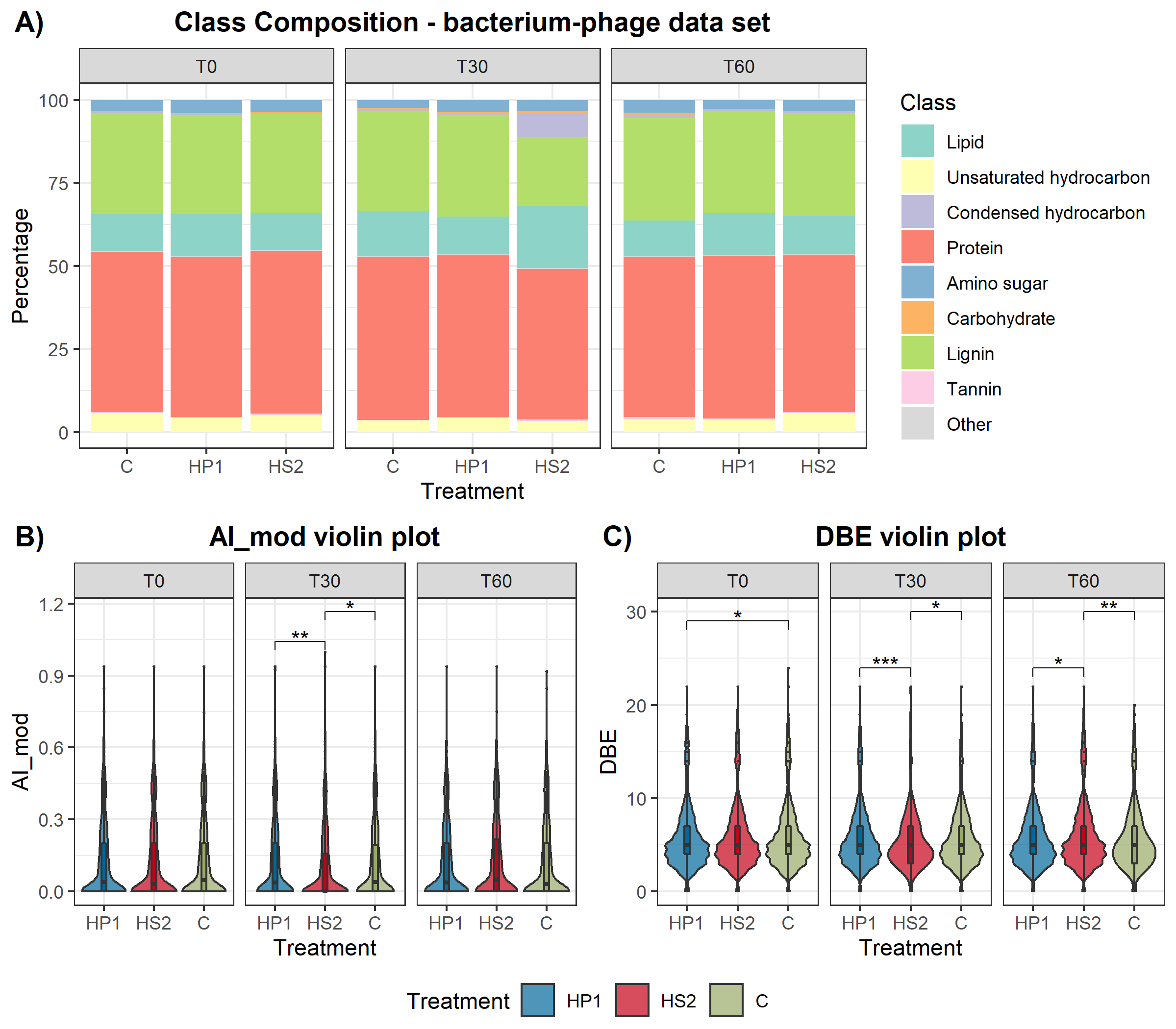

#### Supplementary Figure 3. A) Changes in the molecular class composition of the bacterium-phage exometabolome during the incubation. A reduction in the percentage of lignin-like compounds is observed at 30 minutes after inoculation. B) Violin plot of the changes in the aromatic index reflect the changes in molecular composition, cells infected with the HS2 phage have lower AI_mod_ at 30 minutes after inoculation (Tukey HSD test, p-value < 0.05). C) Violin plot showing that double bond equivalence (DBE) of HS2 is reduced after 30 minutes and remains low until the end of the experiment (Tukey HSD test, p-value < 0.05). For B), C) and D) * (p-value < 0.05), ** (p-value < 0.01), *** (p-value < 0.001), **** (p-value < 0.0001).

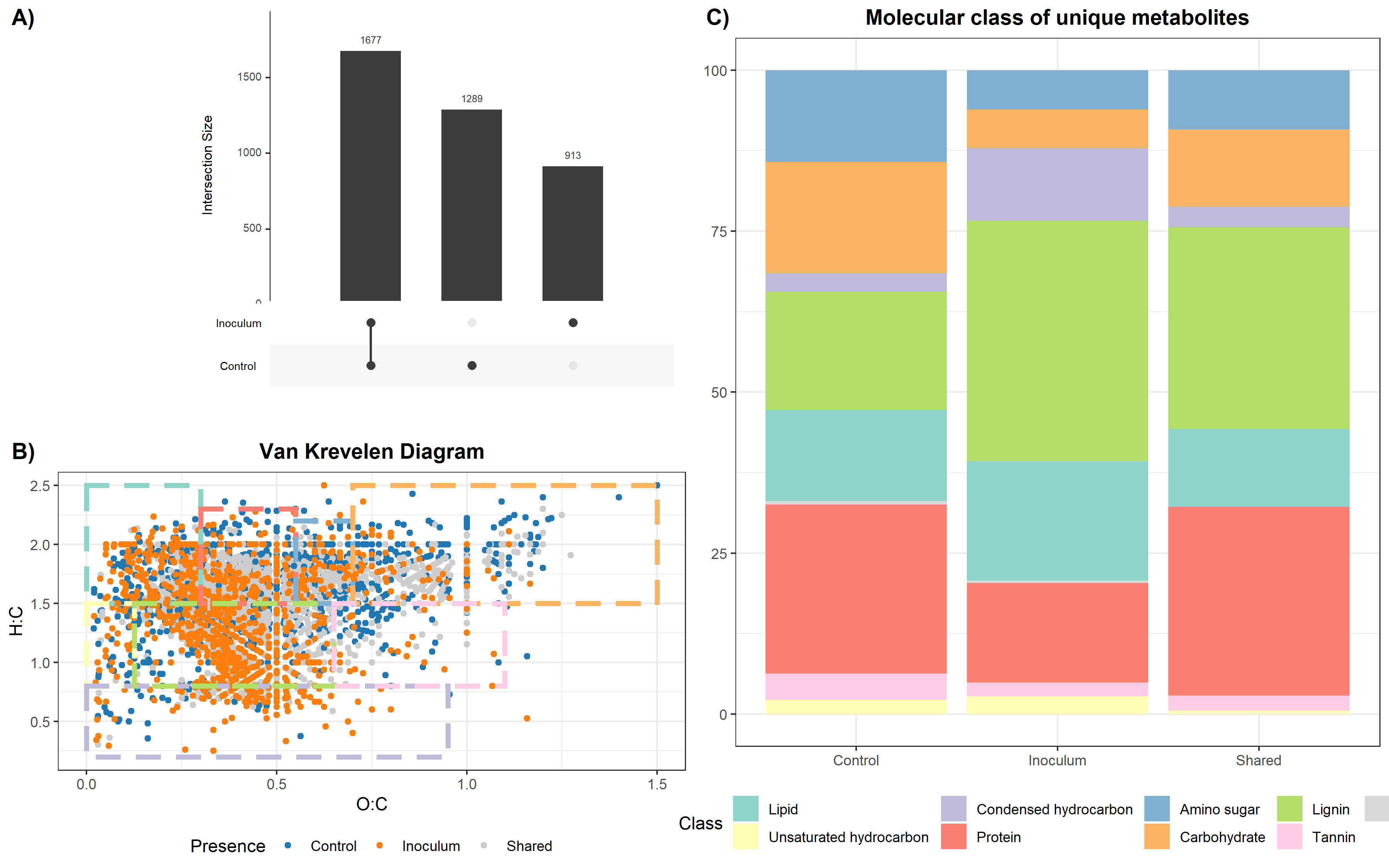

#### Supplementary Figure 4. A) Upset plot showing the number of metabolites that are shared and unique between control and inoculated treatments of the *S. fallax* leachate. B) Van Krevelen diagram showing metabolites that are shared and unique between control and inoculated treatments of the *S. fallax* leachate. C) Molecular composition of the unique metabolites showing that there are unique protein-like, carbohydrate-like, lignin-like and lipid-like metabolites in cells infected with each phage.

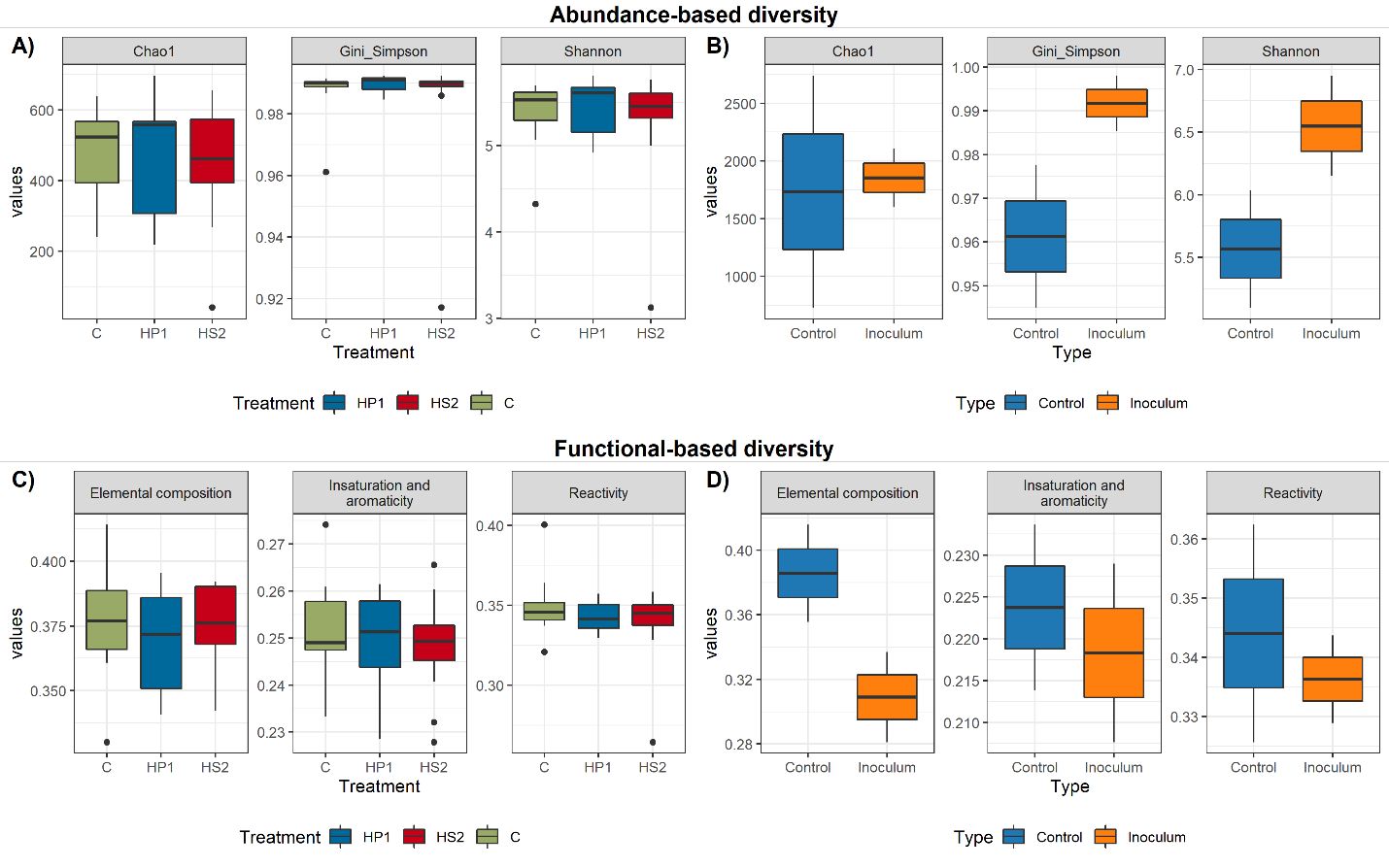

#### Supplementary Figure 5. Results of the chemodiversity analysis of the FTICR-MS data. A) and B) Abundance-based diversity metrics including the Chao1 richness estimator, Gini-Simpson, and Shannon indexes. C) and D) Functional-based diversity using Rao’s quadratic entropy using different traits: Elemental composition is based on the number of elements in each molecular formula. Insaturation and aromaticity uses DBE and AImod as traits. Reactivity uses Gibbs’ free energy as trait.

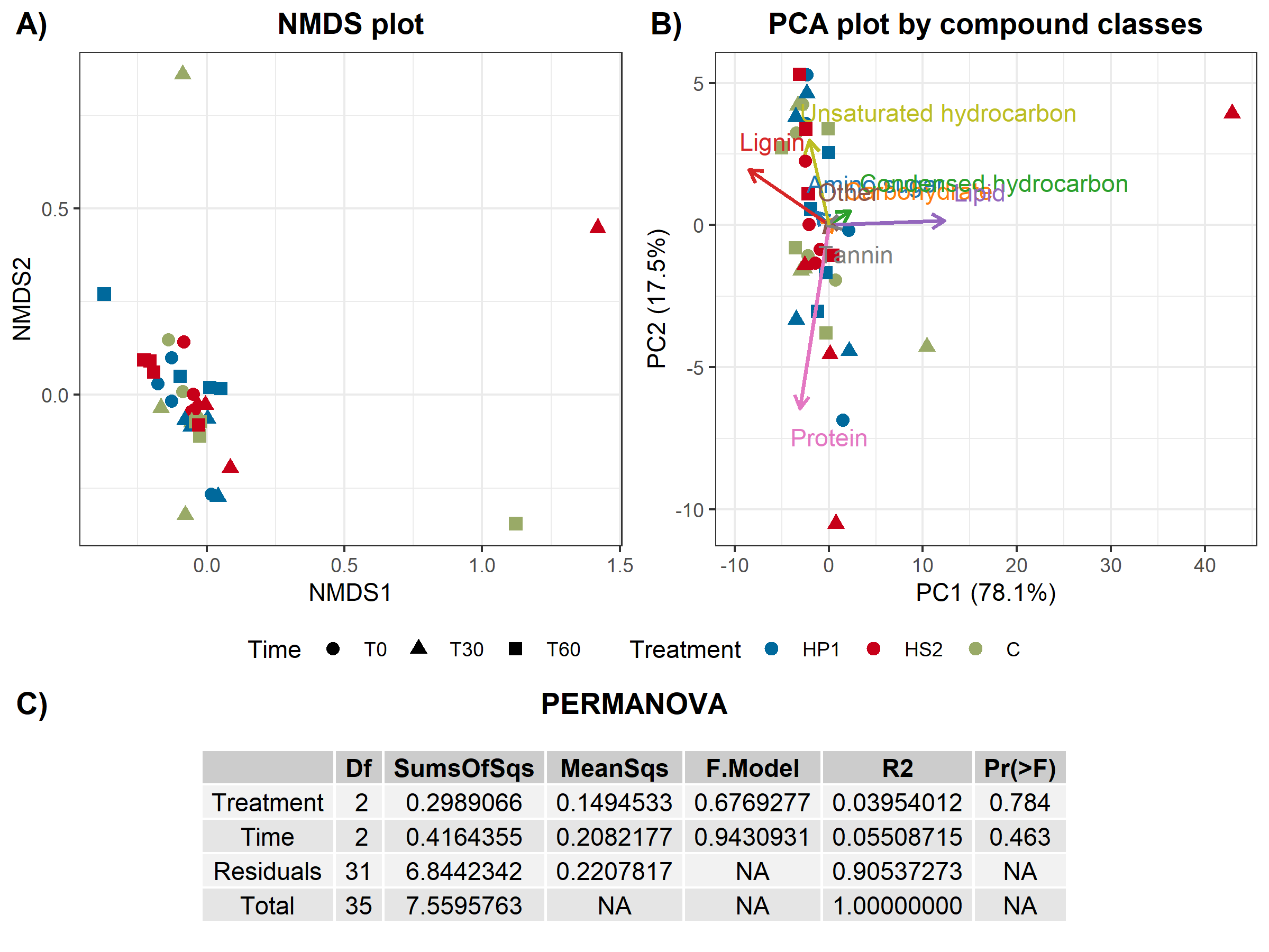

#### Supplementary Figure 6. Multivariate statistical analysis performed by MetaboDirect. A) NMDS plot showed a small clustering of samples based on the content of phosphorus rather than the type of infection, as denoted by the clusters of colored dots B) PCA plots by compounds molecular class. Like the NMDS plot, there was no clustering of the sample neither by phage nor time. C) PERMANOVA result, the last column shows the p-value of the analysis. There was not a significant effect of the phage or the time.
